# Bat and pangolin coronavirus spike glycoprotein structures provide insights into SARS-CoV-2 evolution

**DOI:** 10.1101/2020.09.21.307439

**Authors:** Shuyuan Zhang, Shuyuan Qiao, Jinfang Yu, Jianwei Zeng, Sisi Shan, Jun Lan, Long Tian, Linqi Zhang, Xinquan Wang

**Author notes:** These authors contributed equally to this work (Shuyuan Zhang, Shuyuan Qiao, Jinfang Yu). Correspondence (X.W.).

## Abstract

In recognizing the host cellular receptor and mediating fusion of virus and cell membranes, the spike (S) glycoprotein of coronaviruses is the most critical viral protein for cross-species transmission and infection. Here we determined the cryo-EM structures of the spikes from bat (RaTG13) and pangolin (PCoV_GX) coronaviruses, which are closely related to SARS-CoV-2. All three receptor-binding domains (RBDs) of these two spike trimers are in the “down” conformation, indicating they are more prone to adopt this receptor-binding inactive state. However, we found that the PCoV_GX, but not the RaTG13, spike is comparable to the SARS-CoV-2 spike in binding the human ACE2 receptor and supporting pseudovirus cell entry. Through structure and sequence comparisons, we identified critical residues in the RBD that underlie the different activities of the RaTG13 and PCoV_GX/SARS-CoV-2 spikes and propose that N-linked glycans serve as conformational control elements of the RBD. These results collectively indicate that strong RBD-ACE2 binding and efficient RBD conformational sampling are required for the evolution of SARS-CoV-2 to gain highly efficient infection.

## Introduction

Zoonotic transmission of novel coronaviruses pose a tremendous threat to human health, as evidenced by the emergence of SARS-CoV in 2002-2003, MERS-CoV in 2012 and SARS-CoV-2 since the end of 2019^1-5^. SARS-CoV-2 is responsible for the ongoing global COVID-19 pandemic, which has caused millions of infections and hundreds of thousands of deaths worldwide (https://www.who.int/emergencies/diseases/novel-coronavirus-2019/situation-reports/). Current data suggest that similar to SARS-CoV and MERS-CoV^6^, SARS-CoV-2 likely originated in bats and eventually spread to humans following evolution in intermediate hosts. Coronavirus RaTG13, detected in the horseshoe bat *Rhinolophus affinis* in China’s Yunnan province, was identified as the closest relative of SARS-CoV-2^5^. It shares 96.2% sequence identity with the SARS-CoV-2 genome, reflecting the likely origin of SARS-CoV-2 in bats^5^. Pangolin coronaviruses (PCoV) closely related to SARS-CoV-2 have also been identified in smuggled Malayan pangolins (*Manis javanica*) in China’s Guangxi (GX) and Guangdong (GD) provinces. Analyses of PCoV_GX and PCoV_GD genome sequences indicated a high level of similarity with SARS-CoV-2 (85.5% to 92.4% sequence identity)^7-10^. Whether pangolins are intermediate hosts or a natural reservoir for SARS-CoV-2 remains a topic of debate, and it is still unclear how SARS-COV-2 evolved to infect humans.

The spike (S) glycoprotein of coronaviruses forms a trimer, which plays a critical role in host cell attachment and entry by recognizing its cellular receptor and mediating membrane fusion. Consequently, the spike protein, particularly its receptor-binding domain (RBD), is the principal player in determining the host range of coronaviruses^11^. SARS-CoV-2 utilizes human ACE2 (hACE2) as an essential cellular receptor for infection^5,12^. Complex structural determinations have revealed the interactions between SARS-CoV-2 RBD and hACE2 at an atomic level^13-16^. Cryo-EM studies revealed that the SARS-CoV-2 S trimer, similar to that of SARS-CoV, needs to have at least one RBD in an “up” conformation to bind hACE2^17-23^. Therefore, a spike trimer with all three RBDs “down” is in a receptor-binding inactive state, and the conformational change of at least one RBD from “down” to “up” switches the spike trimer to a receptor-binding active state^18^. The spike and RBD of RaTG13 and SARS-CoV-2 share 97.5% and 89.2% amino acid sequence identity, respectively. Similar to RaTG13, PCoV_GX (GenBank: QIA48614.1) shares 92.3% and 86.7% amino acid sequence identity with the SARS-CoV-2 spike and RBD. In contrast, PCoV_GD (GenBank: QLR06867.1) and SARS-CoV-2 have greater amino acid sequence identity in the RBD (96.9%) than in the spike protein (89.6%). Consistently, the RBD of PCoV_GD has demonstrated stronger binding to hACE2 than the RBD of RaTG13, and hACE2 also supported more efficient cell entry of PCoV_GD than RaTG13 pseudoviruses^24^. Data have not been reported regarding the binding of PCoV_GX spike and its RBD to hACE2 or whether hACE2 supports PCoV_GX pseudovirus cell entry.

Here we report the cryo-EM structures of RaTG13 and PCoV_GX spikes at 2.48 Å and 2.93 Å resolution, respectively. These two spikes have all three RBDs in the “down” conformation. Our structural comparisons of RaTG13, PCoV_GX and SARS-CoV-2 S proteins, coupled with functional data on hACE2 binding and pseudovirus cell entry, provide important insights into the evolution and cross-transmission of SARS-CoV-2.

## Results

### Protein expression and structure determination

The cDNAs encoding the PCoV_GX (GenBank: QIA48614.1) and RaTG13 (GenBank: QHR63300.2) spike proteins were synthesized with codon optimization for recombinant expression. The PCoV_GX ectodomain (residues 1-1205) was cloned into the pCAG vector and the RaTG13 ectodomain (residues 1-1209) into the pFastBac-Dual vector. Both constructs include a C-terminal foldon tag for trimerization, a Strep tag for purification, and the ‘2P’ mutations for protein stabilization (K980P and V981P for PCoV_GX; K982P and V983P for RaTG13). After purification of PCoV_GX spike from FreeStyle 293-F cells and that of RaTG13 from Hi5 insect cells, both proteins existed as heavy glycosylated homotrimers with no cleavage into the S1 and S2 subunits by endogenous proteases (Fig. S1). Cryo-EM images were recorded using a FEI Titan Krios microscope operating at 300 KV with a Gatan K3 Summit direct electron detector. For the PCoV_GX and RaTG13 spike trimers, ~700,000 and ~450,000 particles, respectively, were subjected to 2D classification, and a total of 263,842 and 99,241 particles were selected and subjected to 3D refinement with C3 symmetry to generate density maps (Fig. S2). The overall density maps were solved to 2.48 Å for the PCoV_GX spike and 2.93 Å for the RaTG13 spike (gold-standard Fourier shell correlation = 0.143) (Fig. S3). The atomic-resolution density maps enabled us to build nearly all residues of the PCoV_GX spike (residues 14-1138) with 84 N-linked glycans (Fig. S4). The refined RaTG13 spike model contains residues 14-1133 with seven breaks (residues 19-23, 67-80, 144-156, 176-186, 243-264, 677-685 and 824-830) and 54 N-linked glycans (Fig. S4). Data collection and refinement statistics for these two structures are listed in Table S1.

### Overall structures of RaTG13 and PCoV_GX spikes

The overall structures of homotrimeric RaTG13 and PCoV_GX spikes resemble the previously reported pre-fusion structures of coronavirus spikes (Fig. 1A). Both spikes have a mushroom-like shape (~150 Å in height and ~120 Å in width), consisting of a cap mainly formed by β-strands and a stalk mainly formed by α-helices (Fig. 1A). Like other coronaviruses, the RaTG13 and PCoV_GX spike monomers are composed of the S1 and S2 subunits with a protease cleavage site between them (Fig. 1B,1C). The structural components of the spike include the N-terminal domain (NTD), RBD (also called the C-terminal domain, CTD), subdomain 1 (SD1) and subdomain 2 (SD2) in the S1 subunit; and the upstream helix (UH), fusion peptide (FP), connecting region (CR), heptad repeat 1 (HR1), central helix (CH), β-hairpin (BH), subdomain 3 (SD3) and heptad repeat 2 (HR2) in the S2 subunit (Fig. 1D, Fig. S5).

**Fig.1.**
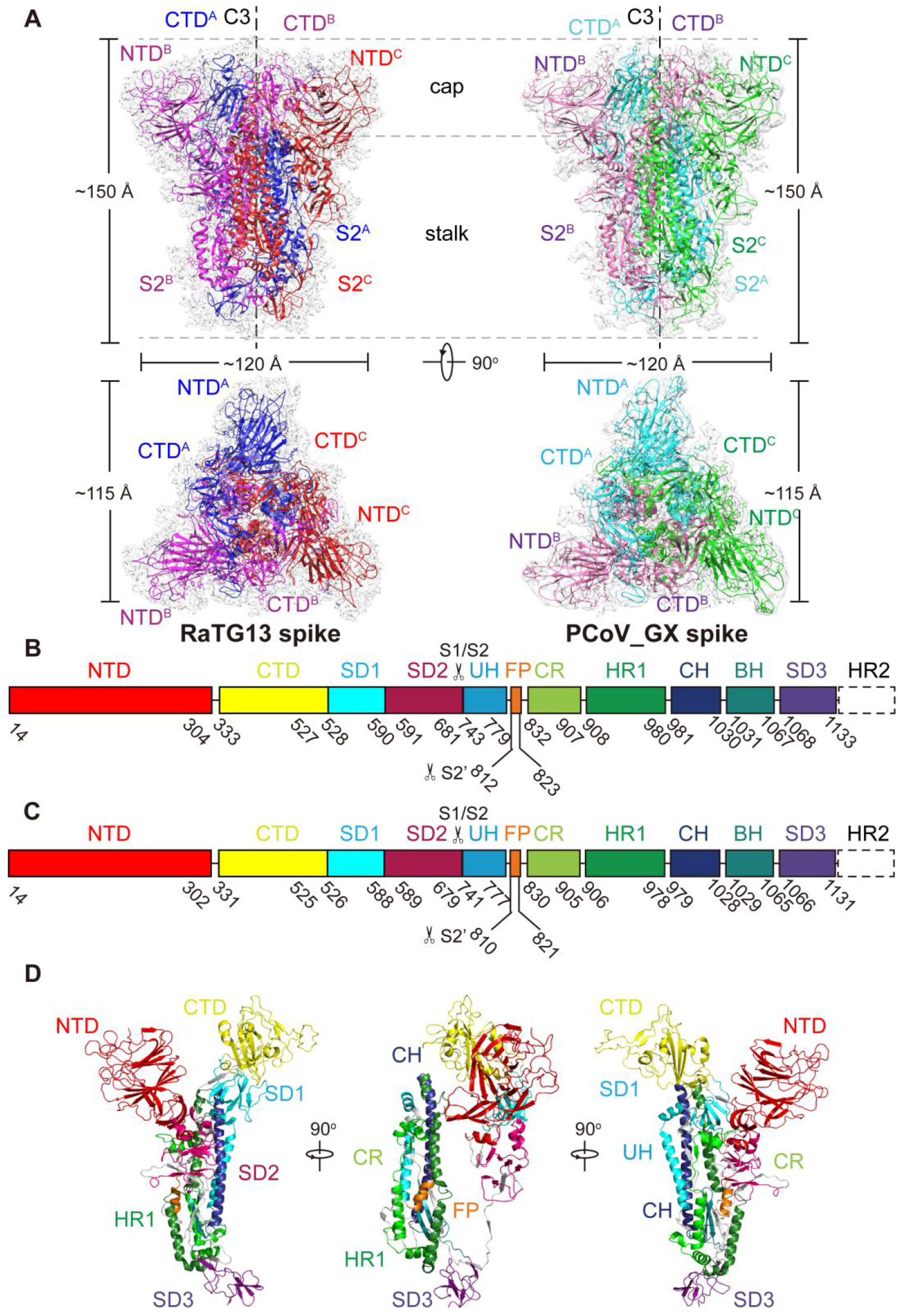
Overall structures of the RaTG13 and PCoV_GX spike glycoproteins. **(A)** Overall structures of RaTG13 and PCoV_GX spike glycoproteins shown in side view (upper panel) and top view (lower panel). Three monomers of the RaTG13 spike are colored magenta, red, and blue, respectively; three monomers of the PCoV_GX spike are colored hot pink, green and cyan, respectively. The cryo-EM maps are shown as a semitransparent surface. The trigonal axes are shown as black dashed lines. Visible segments of each monomer are labeled accordingly. The cap and stalk parts are partitioned by gray dashed lines. **(B)** Schematic representation of the RaTG13 spike monomer structural domains. The domains of RaTG13 are shown as boxes with the width related to the length of the amino acid sequence. The start and end amino acids of each segment are labeled. The position of the S1/S2 and S2’ cleavage sites are indicated by scissors. NTD, N-terminal domain; CTD, C-terminal domain; SD1, subdomain 1; SD2, subdomain 2; UH, upstream helix; FP, fusion peptide; CR, connecting region; HR1, heptad repeat 1; CH, central helix; BH, β-hairpin; SD3, subdomain 3. **(C)** Schematic representation of the PCoV_GX spike monomer structural domains. The abbreviations of elements are the same as in **B. (D)** Cartoon diagrams depicting three orientations of the spike monomer colored as in **B** and **C**. As the RaTG13 and PCoV_GX spike monomers have extremely similar structures, thus only the RaTG13 spike monomer was used to show the detailed architecture.

RaTG13 and PCoV_GX spikes have the typical β-coronavirus structural features^25^. Their NTDs have a core consisting of three β-sheets plus one helix and a ceiling structure above the core (Fig. S6). Three conserved disulfide bonds that are found in other β-coronavirus NTDs are also present in the NTDs of RaTG13 (15C-136C, 131C-166C and 291C-301C) and PCoV_GX (15C-134C, 129C-164C and 289C-299C) (Fig. S6). The RaTG13 and PCoV_GX RBDs adopt an architecture similar to that of other β-coronavirus RBDs, with a β sheet core and an inserted loop called a receptor-binding motif (RBM) (Fig. 2A). Detailed structural descriptions and comparisons of these two RBDs are presented in the next section. The remaining SD1 and SD2 domains in the S1 subunit and the S2 subunits of RaTG13 and PCoV_GX are also structurally conserved and similar to those of SARS-CoV-2.

**Fig.2.**
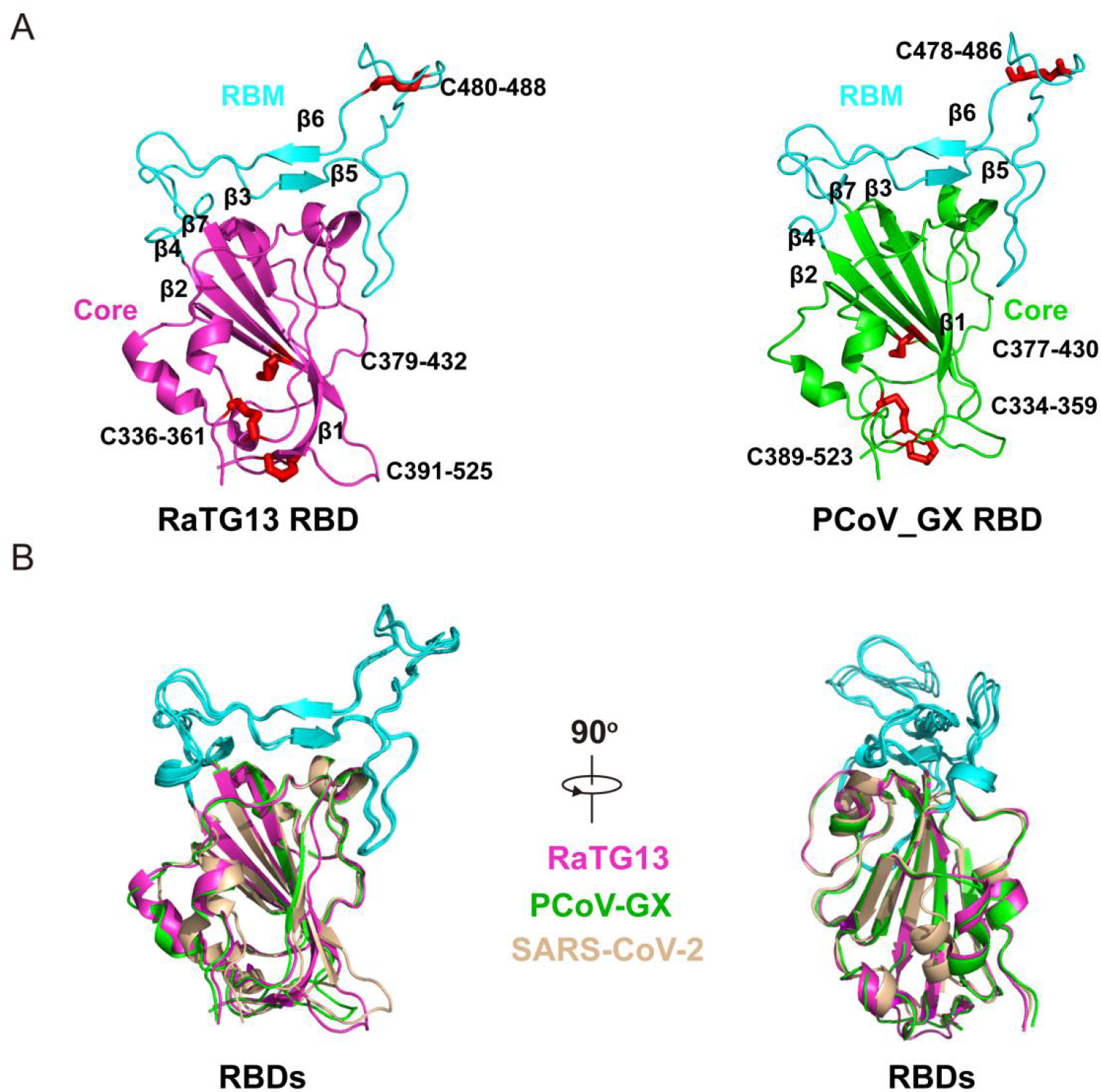
RBD structures of the RaTG13 and PCoV_GX spike proteins. **(A)** The RaTG13 and PCoV_GX RBDs are shown in side view. The RaTG13 RBD core is colored in magenta and the RBM in cyan; the PCoV_GX RBD core is colored in green and the RBM in cyan. Disulfide bonds are shown as red sticks with residues labeled. **(B)** Structural alignment of the RaTG13, PCoV_GX and SARS-CoV-2 (PDB ID:6M0J; core colored in wheat) RBDs. Aligned structures are shown in two orientations.

### Structures and hACE2 binding of the RBDs

Most β-coronaviruses utilize the RBD to specifically bind the host receptor. Compared to other structural components in the spike, the RBD harbors the most sequence and structure variations across different β-coronaviruses and thus has important implications for viral evolution and cross-species transmission. The RaTG13 and PCoV_GX RBD cores are comprised of a twisted five-stranded antiparallel β sheet (β1, β2, β3, β4 and β7) with connecting loops and helices (Fig. 2A). The RBM, a long loop with two short β strands (β5 and β6), is inserted between the β4 and β7 strands (Fig. 2A). Besides three disulfide bonds in the core (336C-361C, 379C-432C and 391C-525C in RaTG13; 334C-359C, 377C-430C and 389C-523C in PCoV_GX) that stabilize the β sheet, the RaTG13 and PCoV_GX RBDs also have an additional disulfide bond (480C-488C in RaTG13 and 478C-486C in PCoV_GX) that connects the loop at the distal end of the RBM (Fig. 2A). The overall structures of the RaTG13/PCoV_GX and SARS-CoV-2 RBDs are highly similar (Fig. 2B). The rmsd for aligned Cα atoms is 0.91 Å between the RaTG13 and SARS-CoV-2 RBDs and 0.59 Å between the PCoV_GX and SARS-CoV-2 RBDs.

We measured the binding affinities of hACE2 with the RBDs of RaTG13, PCoV_GX and SARS-CoV-2 using surface plasmon resonance (SPR). The PCoV_GX and SARS-CoV-2 RBDs bound to hACE2 with comparable affinities of 2.7 nM and 3.9 nM, respectively. However, the RaTG13 RBD bound to hACE2 with a much weaker affinity of 216 nM (Fig. 3A). Sequence comparisons showed that both the RaTG13 and PCoV_GX RBMs share 75.3% amino acid sequence identity with the RBM of SARS-CoV-2. Of the 16 residues in the SARS-CoV-2 RBM involved with ACE2 binding, ten are conserved in RaTG13, PCoV_GX and SARS-CoV-2 (Fig. 3B). The other six SARS-CoV-2 residues that are not conserved in both RatG13 and PCoV_GX are Y449, F486, Q493, Q498, N501 and Y505 (Fig. 3B). Except for F486, which is replaced by a leucine in RaTG13 and PCoV_GX, these residues (Y449, Q493, Q498, N501 and Y505) in the SARS-CoV-2 RBM form a patch that has significant hydrophilic interactions with hACE2 (Fig. 3B, Fig. S7). Of these five positions, SARS-CoV-2 Y449 forms two hydrogen bonds with hACE2 D38 and Q42 upon binding. This tyrosine is conserved in the PCoV_GX RBD but is replaced by a phenylalanine in the RaTG13 RBD, which would disrupt the hydrogen-bonding interactions (Fig. 3C, Fig. S7). Similarly, SARS-CoV-2 Y505 forms two hydrogen bonds with hACE2 E37 and R393. This residue is conserved in PCoV_GX RBD, perhaps having a similar effect in facilitating hACE2 binding, whereas the histidine found at this site in the RaTG13 RBD would alter interactions with hACE2 (Fig. 3C, Fig. S7). We therefore propose that Y449 and Y505 are two of the principal sites that contribute to the weaker binding of the RaTG13 RBD with hACE2 compared to that of the RBDs of PCoV_GX and SARS-CoV-2.

**Fig.3.**
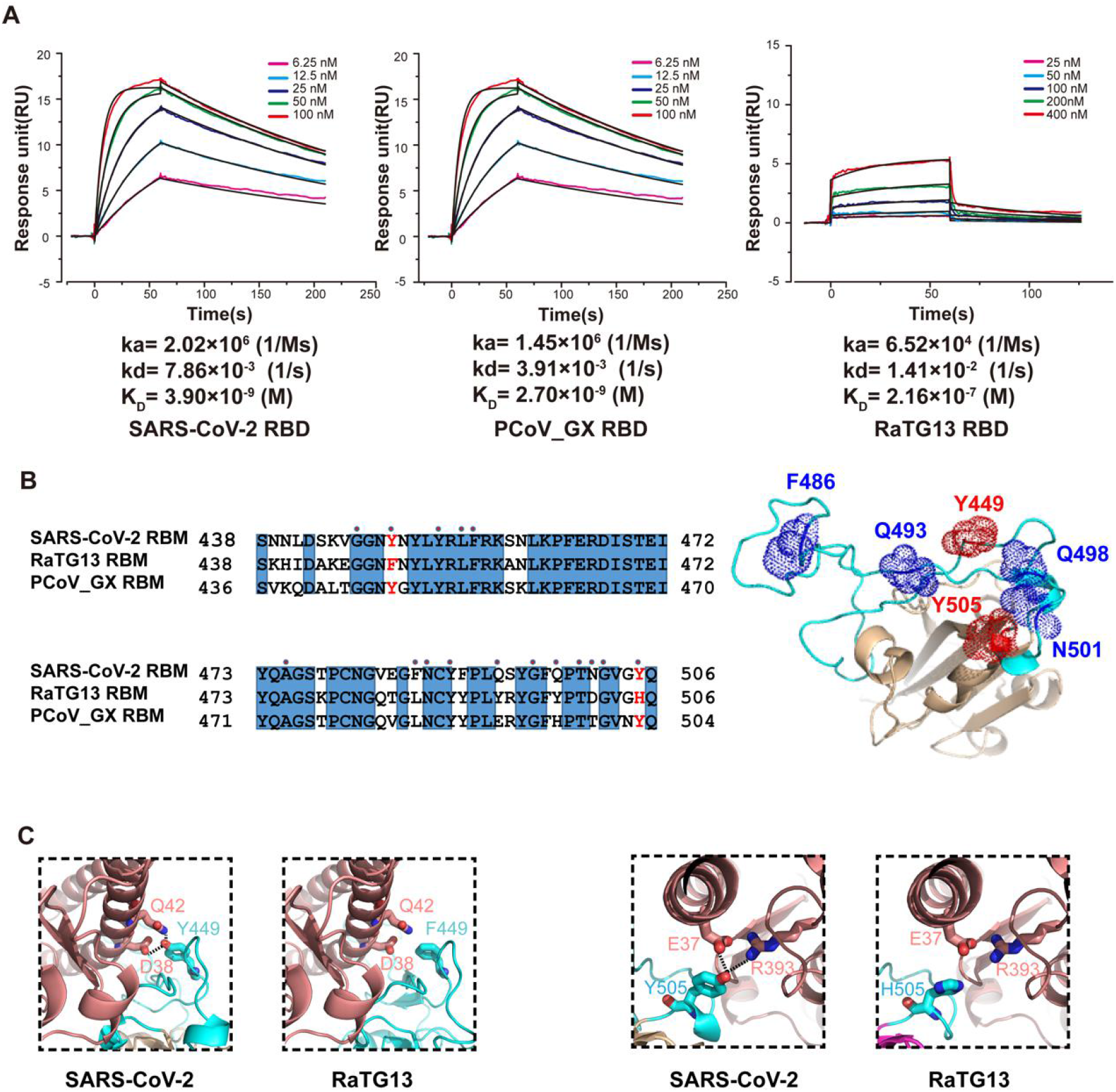
The relationship between binding affinity and sequence variation in different spike proteins. **(A)** Binding curves of immobilized human ACE2 with the SARS-CoV-2 (left panel), PCoV_GX (middle panel) or RaTG13 (right panel) RBD. Data are shown as different colored lines and the best fit of the data to a 1:1 binding model is shown in black. **(B)** Sequence alignment of the RBMs from the SARS-CoV-2, PCoV_GX and RaTG13 spike proteins (left panel). Residues Y449 and Y505 in the SARS-CoV-2 RBM and the corresponding residues in the RaTG13 and PCoV_GX RBMs are marked in red. The RBD of SARS-CoV-2 (PDB ID:6M0J) shown as a cartoon (right panel). Residues in the SARS-CoV-2 RBM that contact hACE2 are indicated by red dots. Residues 486, 493, 498 and 501 in the RBM of SARS-CoV-2 are shown as blue dots. **(C)** Principal residues at the SARS-CoV-2 RBD–hACE2 (PDB ID:6M0J) and RaTG13 RBD–hACE2 interfaces. Hydrogen bonds between SARS-CoV-2 Y449 and hACE2 D38 and Q42 would be abolished after Y to F mutation in the RaTG13 RBM (two leftmost panel). Hydrogen bonds between SARS-CoV-2 Y505 and hACE2 E37 and R393 would be abolished after Y to H mutation in the RaTG13 RBM (two rightmost panels).

### Conformations and hACE2 binding of the RaTG13 and PCoV_GX spikes

As described above, at least one of the RBDs in the SARS-CoV-2 spike trimer must adopt an “up” conformation in order to bind hACE2. By cryo-EM, we only captured conformational states of RaTG13 and PCoV_GX spikes with all three RBDs in the “down” position. Structures of SARS-CoV-2 spike trimer with all three RBDs in the “down” conformation were previously determined (PDB IDs: 6VXX and 6ZGE)^19,21^. In 6VXX, one of the SARS-CoV-2 RBDs exhibits contacts with 11 residues and the N165/N234-linked glycans from the counter-clockwise monomer and 5 residues from the clockwise monomer (Fig. 4A and 4B, Table S2). In 6ZGE, the three “down” RBDs are more compactly packed, with one RBD having interactions with 35 residues and the N165/N234-linked glycans of the two neighboring monomers (Fig. 4A and 4B, Table S2). In the RaTG13 spike, one RBD contacts 27 residues and the N165/N234/N370-linked glycans from the counter-clockwise monomer, and 13 residues from the clockwise monomer with a distance cutoff of 4.0 Å (Fig. 4A and 4B, Table S2). A nearly identical packing of RBDs was also observed in a recently reported RaTG13 spike structure (PDB ID:6ZGF)^21^. In the PCoV_GX spike, the number of RBD-contacting residues is 27 from the counter-clockwise monomer and 16 from the clockwise monomer. The N163/N232/N368-linked glycans from the counter-clockwise monomer are also involved in contact with the RBD (Fig. 4B, Table S2). Therefore, regarding RBD packing, the RaTG13 and PCoV_GX spikes are more similar to the SARS-CoV-2 spike structure 6ZGE than 6VXX.

**Fig.4.**
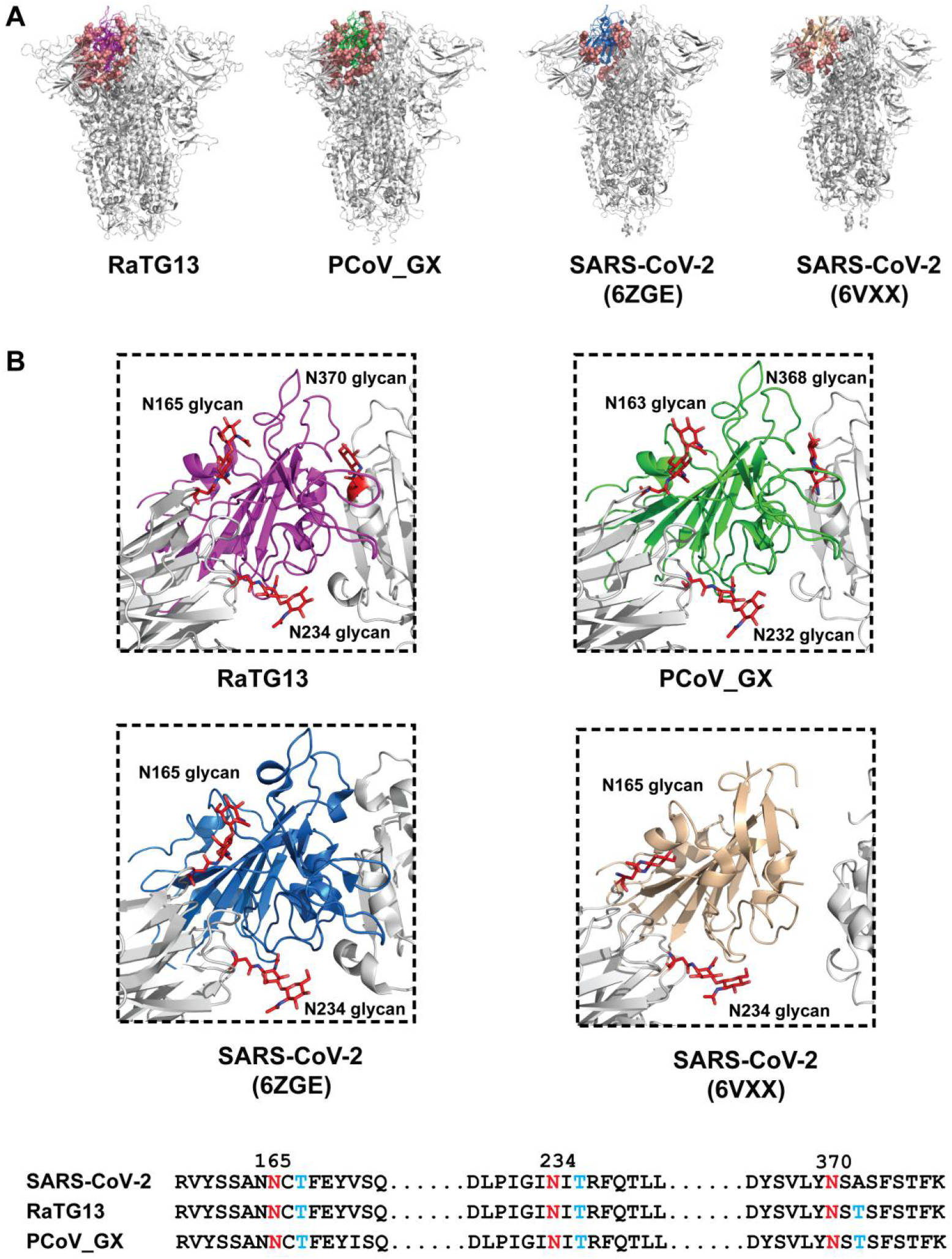
The residues and glycans interacting with one RBD of the different spikes. **(A)** The residues and glycans interacting with one RBD are shown as salmon spheres. The RaTG13 RBD is colored in magenta, PCoV_GX RBD in green, SARS-CoV-2 (PDB ID: 6VXX) RBD in wheat, and SARS-CoV-2 (PDB ID: 6ZGE) RBD in marine; remaining regions shown in gray. **(B)** Detailed structures of the RBD-glycans interface are shown. The RaTG13, PCoV_GX and SARS-CoV-2 (PDB ID: 6ZGE/6VXX) RBDs are colored the same as in **A**. Glycans are shown as red sticks and Asn-linked glycans are labeled. Sequence alignment of the SARS-CoV-2, RaTG13 and PCoV_GX RBD-interacting glycosylation sites is shown in the bottom panel. Some sequences between the three sites are omitted and indicated by black dots. Amino acid positions of asparagines are indicated above the sequences according to SARS-CoV-2. Asparagines (N) are colored red and threonines (T) are colored blue.

Unlike in our study, the cryo-EM studies which determined the structures of 6ZGE and 6VXX did capture the SARS-CoV-2 spikes adopting a more loose state with one “up” RBD. Our observations of all three RBDs only in the “down” position in the RaTG13 and PCoV_GX spikes suggests they are more prone to adopt the receptor-binding inactive state. Considering that the number of protein-protein interactions around the “down” RBD is nearly the same among the RaTG13, PCoV_GX and SARS-CoV-2 (6ZGE) spikes, glycans may play an important role in how efficiently the RBD can sample different conformations. Of note, we observed contacts between the RBD and three neighboring N-linked glycans, spatially positioned at three vertices of a triangle, in the RaTG13 and PCoV_GX spikes (Fig. 4B). Although the SARS-CoV-2 spike also has three asparagine residues (N165, N234 and N370) at these same positons, N370 is not a glycosylation site in the SARS-CoV-2 spike and thus glycans contacting the RBD are not observed at this positon (Fig. 4B).

To further our findings, we also measured the binding affinities of hACE2 with the spikes of RaTG13, PCoV_GX and SARS-CoV-2. Interestingly, we found that despite exhibiting only a receptor-binding inactive conformation in the cryo-EM images, the PCoV_GX spike bound hACE2 with an affinity of 130 nM, comparable to the 105 nM affinity of the SARS-CoV-2 spike (Fig. 5A). The binding of RaTG13 spike to hACE2 was weaker, with an affinity of 600 nM. We tested the entry of RaTG13, PCoV_GX and SARS-CoV-2 pseudoviruses into HEK293 cells expressing hACE2. Consistently, the PCoV_GX and SARS-CoV-2 pseudoviruses had comparable entry efficiency, whereas the RaTG13 pseudovirus exhibited little to no entry (Fig. 5B).

**Fig.5.**
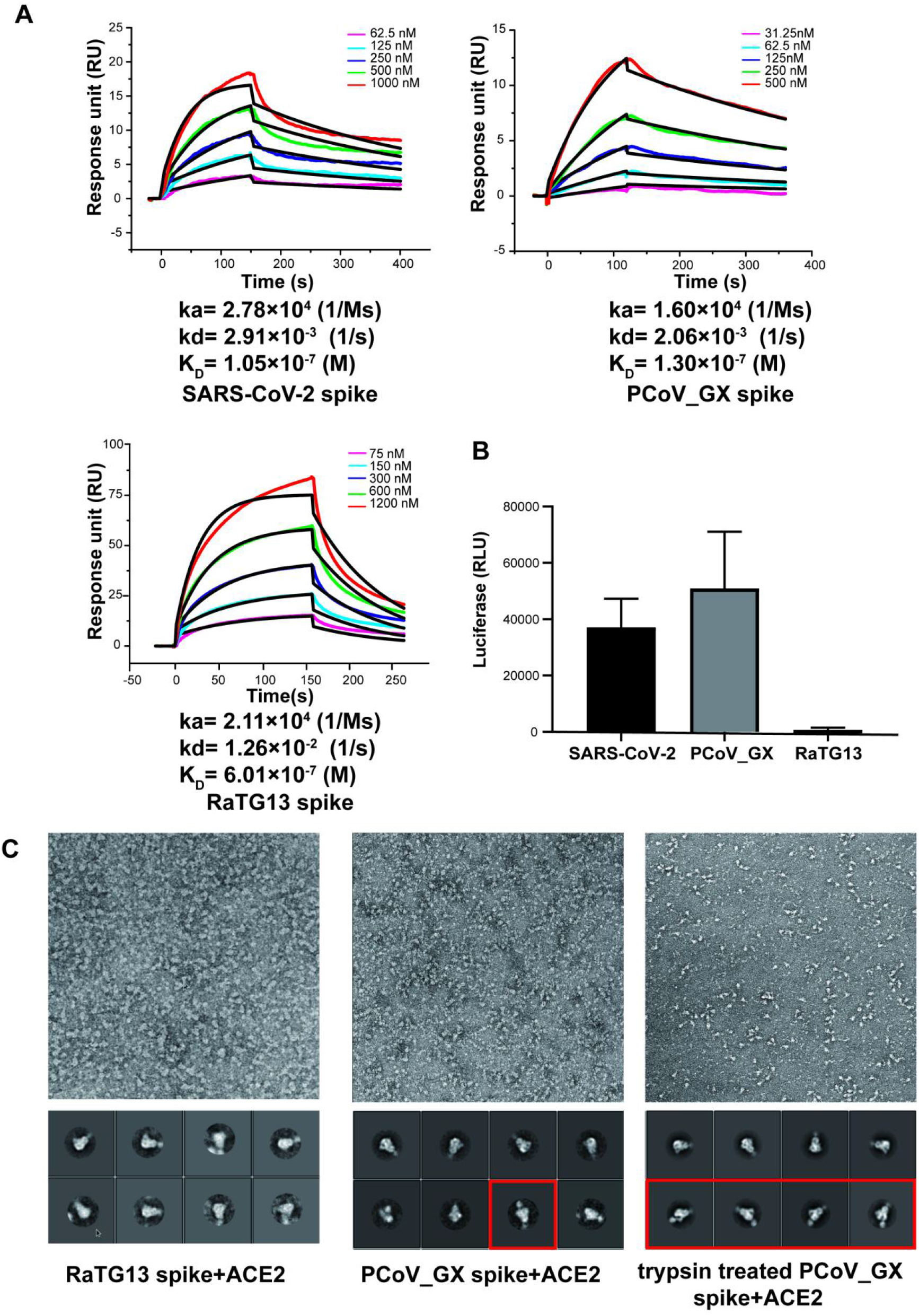
Binding affinities and cell entry of the different spikes. **(A)** Binding curves of immobilized hACE2 with the SARS-CoV-2, PCoV_GX or RaTG13 spike. Data are shown as different colored lines and the best fit of the data to a 1:1 binding model is shown in black. **(B)** The cell entry efficiencies of pseudotyped viruses as measured by luciferase activity. SARS-CoV-2, PCoV_GX and RaTG13 pseudotyped viruses were used to infect hACE2-transfected HEK293 cells. **(C)** The representative micrographs and 2D classification results of negative-staining EM. Both spikes were incubated with 4-fold molar ratio of hACE2. The red box shows the complex of the PCoV_GX spike with hACE2.

To capture the hACE2-bound state of the RaTG13 and PCoV_GX spikes, we mixed spike and ACE2 at a 1:4 molar ratio and performed negative-staining EM. The 2D classification did not show particles with bound hACE2 for the RaTG13 spike, but ~9% of the PCoV_GX particles were bound to hACE2. After treating the PCoV_GX spike with trypsin for 2 hours, the ratio of hACE2-bound particles increased to 20% (Fig. 5C). These results further support that the conformational switch of the spike is a dynamic equilibrium process and that binding of hACE2 would capture the spike with “up” RBDs and shift the process towards more spikes ready for receptor binding and membrane fusion.

## Discussion

Coronavirus spike glycoproteins recognize their host cellular receptor and mediate membrane fusion for entry, thereby functioning as the most critical coronavirus protein in determining viral evolution and cross-species transmission. In this study, cryo-EM structures of RaTG13 and PCoV_GX spikes were determined at atomic resolution. Our comparisons of the structures of the RaTG13, PCoV_GX and SARS-COV-2 spikes, the strength of their hACE2-binding, and their efficiency in facilitating pseudovirus cell entry provide important insights into the evolution and cross-species transmission of SARS-CoV-2.

Our structural determinations of the RaTG13 and PCoV_GX spikes showed that the RBDs of these two coronaviruses are highly similar to that of SARS-CoV-2. However, our SPR experiments showed that only PCoV_GX RBD exhibited a hACE2-binding affinity comparable to SARS-CoV-2 RBD, whereas RaTG13 RBD demonstrated far weaker binding. Sequence alignments showed that variation at six residues (SARS-CoV-2 Y449, F486, Q493, Q498, N501 and Y505) were responsible for the these differences in hACE2-binding among the RaTG13, PCoV_GX and SARS-CoV2 RBDs. The residues Y449, Q493, Q498, N501 and Y505 are especially important, clustering together to form a patch on the SARS-CoV-2 RBD that has significant interactions with hACE2. We further pinpointed amino acid changes at two positions (Y449 and Y505) only seen in the RaTG13, and not the PCoV_GX, RBD that may account for the weaker binding we observed between hACE2 and the RaTG13 RBD. Our findings and conclusions are supported by recent reports of adapted and remodeled SARS-CoV-2 strains utilized in mouse model studies. Gu et al. reported an adapted SARS-CoV-2 strain with increased infectivity in mice that has a N501Y mutation in the RBD^26^. Dinnon et al. remodeled the SARS-COV-2 RBD at two sites (Q498Y and P499T) to facilitate efficient binding to mouse ACE2, producing a recombinant virus that can effectively utilize mouse ACE2 for entry^27^. These positions are within the patch we observed and suggest their importance in the binding capabilities of the RaTG13, PCoV_GX and SARS-CoV-2 RBDs to human ACE2. We further propose that the patch containing Y449, Q493, Q498, N501 and Y505 plays a critical role in the evolution of the SARS-CoV-2 RBD, promoting especially tight binding to hACE2 and impacting the varying affinities observed between the RBD and ACE2 orthologs in wild and domestic animals^24,28^.

The spikes of diverse coronaviruses infecting humans, mice, swine and other hosts have been structurally determined^25^. Current data show that the spikes of only the highly pathogenic human coronavirus SARS-CoV, MERS-CoV and SARS-CoV-2 have a unique structural feature, with the three RBDs in the trimer adopting “down” or “up” conformations^17,19-21,29^. Structure determination of the spike-receptor complex has provided further confirmation that the “up” conformation is required for receptor binding, indicating that the sampling of “down” and “up” conformations by at least one RBD is a prerequisite for receptor binding^18,22,23^ in addition to specific interactions between the RBD and cellular receptor. In our cryo-EM study, the RaTG13 and PCoV_GX spikes exhibited only a receptor-binding inactive state, with all three RBDs adopting the “down” conformation. Another group had the same conclusion in a recent structural determination of the RaTG13 spike^21^. However, in studies of the SARS-CoV-2 spike, the protein seemed to have two conformations, even in the receptor-binding inactive state, with one having more compact packing of the three “down” RBDs than the other^19,21^. We found that the spikes of RaTG13 and PCoV_GX are more similar to the SARS-CoV-2 spike with tight RBD packing. The molecular basis of the more efficient conformational sampling of the SARS-CoV-2 RBD is still not well understood. We observed three N-linked glycans (SARS-CoV-2 positions: N165, N234 and N370) contact the RBD in the RaTG13 and PCoV_GX spikes, whereas N370 is not a glycosylation site in the SARS-CoV-2 spike. The absence of glycans linked to N370 may contribute to the more flexible RBDs of the SARS-CoV-2 spike. This is also supported by a recent study showing that mutation of SARS-CoV-2 N165 resulted in an increase of “up” RBDs, suggesting that glycans serve as a conformational control element of the RBD^30^. We also cannot exclude other factors, such as the furin site that enables cleavage of the spike protein into the S1 and S2 subunits during biogenesis, may also contribute to the RBD flexibility.

The RaTG13 and PCoV_GX spikes and their RBDs all bound hACE2 in our SPR experiments, although both the RBD and spike of PCoV_GX exhibited higher binding affinities than those of RaTG13. These results suggest that RaTG13 and PCoV_GX spikes can also spontaneously sample “up” RBD, which is essential for hACE2 binding. The reason for not observing these conformations in our cryo-EM study may be due to the ratio of RaTG13 and PCoV_GX spike particles adopting this state being too low. Interestingly, the PCoV_GX spike bound to hACE2 with an affinity comparable to that of the SARS-CoV-2 spike and also had similar efficiency in cell entry. In contrast, the RaTG13 spike was much weaker in binding hACE2 and mediating cell entry. We also confirmed the binding of PCoV_GX spike to hACE2 by negative-staining EM.

Based on all these results, we propose that the tight RBD-hACE2 binding we observed is the most critical factor in determining the varied cell-entry efficiency among RaTG13, PCoV_GX and SARS-COV-2. This and the RBD “down” to “up” conformational change are both required for the evolution of SARS-CoV-2 to gain highly efficient transmission capability.

## Materials and Methods

### Protein expression and purification

The cDNAs encoding the SARS-CoV-2 spike (GenBank: YP_009724390.1), PCoV_GX spike (GenBank: QIA48614.1) and RaTG13 spike (GenBank: QHR63300.2) were synthesized with codons optimized for human expression. The SARS-CoV-2 spike ectodomain (1-1121) and PCoV_GX ectodomain (1-1205) were cloned into the pCAG vector separately, and the RaTG13 spike ectodomain (1-1209) was cloned into the pFastBac-Dual vector (Invitrogen). All the spike constructs included a C-terminal foldon tag for trimerization, a Strep tag for purification and ‘2P’ mutations^31^ (K986P and V987P for SARS-CoV-2, K980P and V981P for PCoV_GX, K982P and V983P for RaTG13).

The human ACE2 extracellular domain (19-615), SARS-CoV-2 RBD (333-527), PCoV_GX RBD (331-524) and RaTG13 RBD (333-526) were inserted into the pFastBac-Dual vector, with an N-terminal gp67 signal peptide for secretion and a C-terminal 6 × his tag for purification.

The SARS-CoV-2 and PCoV_GX spike ectodomains were expressed in FreeStyle 293-F cells. Cell cultures were transfected with 1mg of plasmid per liter of culture at a density of 2×10^6^/ml using polyethylenimine (Sigma). The supernatants were collected 72 hours later. RaTG13 spike, SARS-CoV-2 RBD, PCoV_GX RBD, RaTG13 RBD and ACE2 were produced in Hi5 insect cell using the Bac-to-Bac baculovirus system (Invitrogen). Briefly, the amplified high-titer baculoviruses were used to infect Hi5 cells at a density of 2×10^6^/ml, and the supernatants were harvested after 60 hours. SARS-CoV-2, PCoV_GX and RaTG13 spikes were captured by StrepTactin beads (IBA) and further purified by gel-filtration chromatography using a Superose 6 column (GE Healthcare) with buffer containing 20mM Tris-HCl (pH 8.0) and 150mM NaCl. hACE2, SARS-CoV-2 RBD, PCoV_GX RBD and RaTG13 RBD were purified by sequentially applying Ni-NTA resin (GE Healthcare) to a Superdex 200 column (GE Healthcare) with HBS buffer (10 mM HEPES, pH 7.2, 150 mM NaCl).

### Surface plasmon resonance experiments

Running buffer composed of 10 mM HEPES, pH 7.2, 150 mM NaCl and 0.05% (v/v) Tween-20 was used during the analysis, and all proteins were exchanged to the same buffer. hACE2 was immobilized on a CM5 sensorchip (GE Healthcare) at around 700 response units using Biacore T200 (GE Healthcare). The blank channel of the chip was used as the negative control. Serial dilutions of the SARS-CoV-2, PCoV_GX and RaTG13 spikes and their respective RBDs were flowed through the ACE2 immobilized CM5 chip sequentially. The resulting data were analyzed using Biacore Evaluation Software (GE Healthcare) by fitting to a 1:1 binding model.

### Pseudovirus entry assays

SARS-CoV-2, PCoV_GX and RaTG13 pseudoviruses were generated by co-transfection of human immunodeficiency virus backbones expressing firefly luciferase (pNL43R-E-luciferase) and pcDNA3.1 (Invitrogen) expression vectors encoding the respective spike protein into 293T cells (ATCC). Viral supernatants were collected 48-72 h later. The concentration of the harvested pseudotyped virions was normalized by a p24 ELISA kit (Beijing Quantobio Biotechnology Co., Ltd., China) before infecting hACE2-transfected 293T cells. The infected cells were lysed 24 h after infection and viral entry efficiency was quantified by comparing the luciferase activity among pseudotyped viruses.

### Trypsin treatment of the PCoV_GX and RaTG13 spike glycoproteins

L-(tosylamido-2-phenyl) ethyl chloromethyl ketone (TPCK)-treated trypsin was added to the purified PCoV_GX spike at a mass ratio of 1:100 in HBS buffer and incubated at room temperature for 2 hours. SDS–PAGE was performed to determine that the spikes were fully cleaved into S1 and S2 fragments. The digestion reaction was stopped by applying the mixture to negative staining.

### Negative stain EM

The RaTG13, PCoV_GX and trypsin-cleaved PCoV_GX spikes (0.05mg/ml) were separately mixed with hACE2 on ice for a few minutes at a molar ratio of 1:4, and then deposited onto glow-discharged grids with a continuous carbon layer (Beijing Zhongjingkeyi Technology Co., Ltd.). Excess sample was removed using filter paper after 1 minute of incubation on the grid, then washed twice, incubated with 5 μl of 2% uranyl acetate (UA) solution for another minute, and finally blotted with filter paper. These grids were examined under an FEI Tecnai Spirit electron microscope equipped with an FEI Eagle 4k CCD camera. Images were manually collected at 52,000 magnification with a defocus range between 1.5-1.8 um, corresponding to a pixel size of 2.07 Å. Appropriately, 50 pieces of images were collected for each sample. Image format converting was conducted by EMAN^32^. Particle auto-picking, particle extraction and 2D classification were performed in RELION^33^.

### Cryo-EM sample preparation and data collection

Aliquots of spike ectodomains (4ul, 0.3mg/ml, in buffer containing 20mM Tris-HCl pH 8.0, 150mM NaCl) were applied to glow-discharged holey carbon grids (Quantifoil, Au 300 mesh, R1.2/1.3) and grids with a layer of continuous ultrathin carbon film (Ted Pella, Inc.). The grids were then blotted and plunge-frozen into liquid ethane using an FEI Vitrobot Mark IV.

Images were recorded using FEI Titan Krios microscope operating at 300 kV with a Gatan K3 Summit direct electron detector (Gatan Inc.) at Tsinghua University. The automated software (AutoEMation) was used to collect 3963 movies for PCoV_GX and 1889 movies for RaTG13 at 81,000 magnification at a defocus range between 1.5-1.8 um. Each movie has a total accumulated exposure of 50 e^-^/Å^2^ fractionated in 32 frames of 175 ms exposure. The final image was bin averaged to give a pixel size of 1.0825 Å. Data collection statistics are summarized in Table S1.

### Cryo-EM data processing

Motion Correction (MotionCo2^34^), CTF-estimation (GCTF^35^) and non-templated particle picking (Gautomatch, http://www.mrc-lmb.cam.ac.uk/kzhang/) were automatically executed by TsingTitan.py program. Sequential data processing was carried out on RELION. Initially, ~700,000 particles for PCoV_GX and ~450,000 particles for RaTG13 were subjected to 2D classification. After two or three additional 2D classification, the best selected 474,499 particles for PCoV_GX and 107,274 particles for RaTG13 were applied for initial model and 3D classification.

For PCoV_GX, the best class (397,362 particles) from 3D classification yielded a resolution of 3.14 Å (with C3 symmetry). To improve map density, especially NTD and glycosides, particles were expanded with C3 symmetry, and then subjected to local search classification. The particles of best class from local search classification were further applied to CTF refinement with C3 symmetry and Bayesian polishing, which improved the resolution to 2.71 Å and 2.48 Å, respectively. Meanwhile, the selected particles were subjected to focused classification with an adapted mask on NTD, and then further applied to 3D-refinement, CTF refinement and Bayesian polishing to reach a resolution of 3.64 Å. Additional 3D classification and Bayesian polishing resulted in the NTD map at a resolution of 3.68 Å with better quality. Three copies of NTD maps were fitted onto the whole structure map using Chimera, then combined together using PHENIX combine_focused_maps.

For RaTG13, the best class (99,241 particles) from 3D classification were subjected to 3D auto-refine with C3 symmetry to generate a density map with a resolution of 2.93 Å.

The reported resolutions were estimated with a gold-standard Fourier shell correlation (FSC) cutoff of 0.143 criterion. Local resolution variations were estimated using ResMap^36^. Data processing statistics are summarized in Table S1.

### Model building and refinement

The initial model of PCoV_GX and RaTG13 spikes were generated using the SWISS-MODEL^37^ and fit into the map using UCSF Chimera^38^. Manual model rebuilding was carried out using Coot^39^ and refined with PHENIX real-space refinement^40^. The quality of the final model was analyzed with Molprobity^41^ and EMRinger^42^. The validation statistics of the structural models are summarized in Table S1.

## Acknowledgements

We thank the Tsinghua University Branch of China National Center for Protein Sciences (Beijing) for the cryo-EM facility and the computational facility support, and J. Lei, X. Li, F. Yang, J. Wen and S. Zhang for technical support. This work was supported by funds from the National Key Plan for Scientific Research and Development of China (2016YFD0500307 and 2020YFC0845900) and Tsinghua University Spring Breeze Fund (2020Z99CFY031).

## Author contributions

S.Z. and J.Y. carried out protein expression, purification, electron microscopy sample preparation and data collection with assistance from L.T. S.Z. and S.Q. performed image processing and model building with the help of J.Y. and J.Z. S.Z. and S.Q. performed SPR experiments with assistance from J.L. S.S. conducted pseudovirus entry assays. X.W. and J.Y. conceived, designed and directed the study. X.W., S.Z., S.Q., J.Y.and L.Z. analyzed the structures, made the figures and wrote the manuscript.

## Conflict of interest statement

The authors declare no competing interests

## Notes

### Competing Interest Statement

The authors have declared no competing interest.

