## Supplementary for "Bat and pangolin coronavirus spike glycoprotein structures provide insights into SARS-CoV-2 evolution"

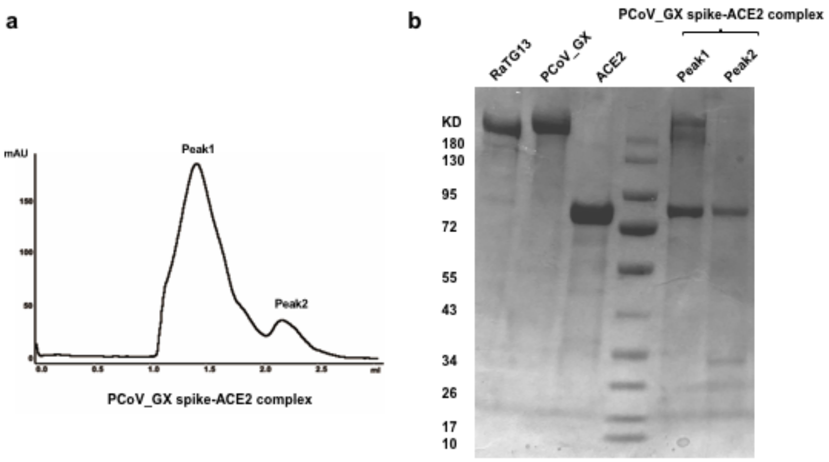


**Supplementary Fig. 1 Purification of the PCoV_GX spike-hACE2 complex. (a)** Size-exclusion chromatography elution profile of the PCoV_GX spike-hACE2 complex. Purified PCoV_GX spike was mixed with hACE2 at a molar ratio of 1:4 before applying to gel-filtration. **(b)** SDS–PAGE analysis of the gel-filtration elutions. Bands on the left side of marker are purified proteins (RaTG13 spike, PCoV_GX spike and hACE2) and bands on the right side are complex. Peak 1, PCoV_GX spike-hACE2 complex; Peak 2, excessive hACE2 .

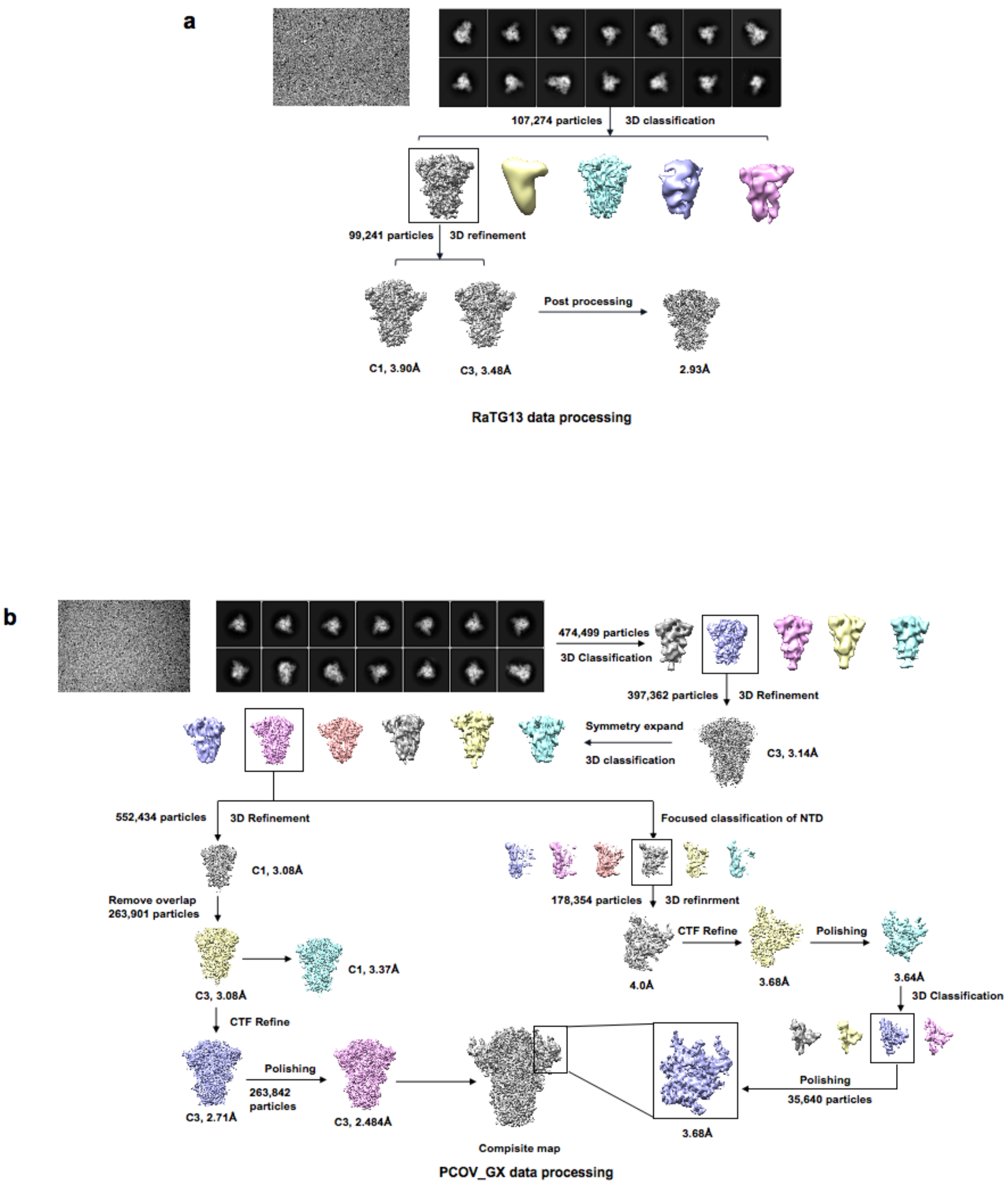


**Supplementary Fig. 2 Cryo-EM data processing workflow.** **(a)** Processing workflow of the RaTG13 spike Cryo-EM data. **(b)** Processing workflow of the PCoV_GX spike Cryo-EM data.

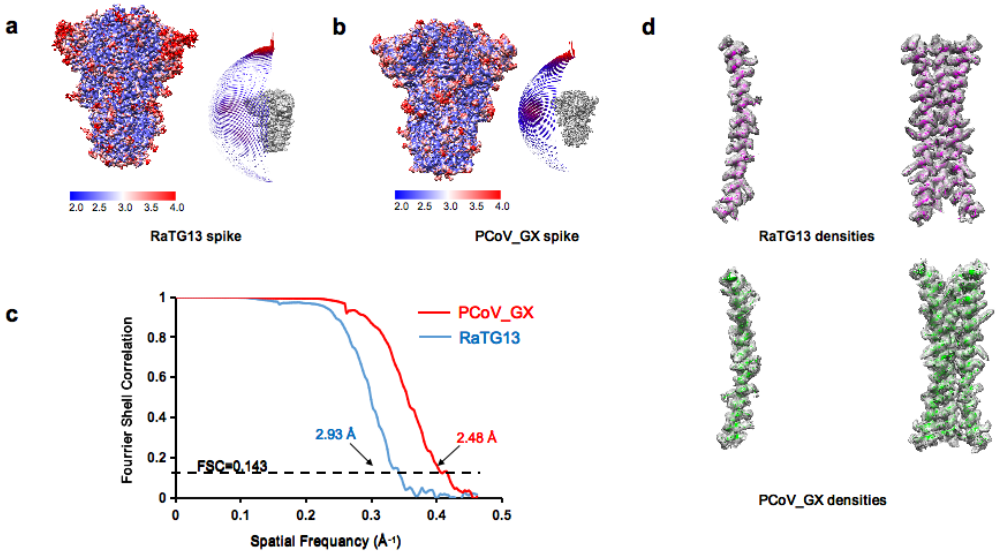


**Supplementary Fig. 3 Cryo-EM structure validations.** Local resolution map (left panel) and particle orientation distribution (right panel) of the RaTG13 spike **(a)** and the PCoV_GX spike **(b)**. A color scale at the bottom of each local resolution map indicates resolution (2 Å to 4 Å). The angular distribution of the particles in each right panel is represented in the three-fold symmetric spike map. **(c)** Gold-standard Fourier Shell Correlation (FSC) curves of the density maps with C3 symmetry. The final resolution of the RaTG13 spike is 2.93 Å. The final resolution of the PCoV_GX spike is 2.48 Å. The 0.143 cut-off value is indicated by a black dashed line. **(d)** Densities from the S2 regions (central helix) of the C3-refined RaTG13 spike (upper panel) and PCoV_GX spike (lower panel) structures. The map is contoured at 2.5 RMS to show the density.

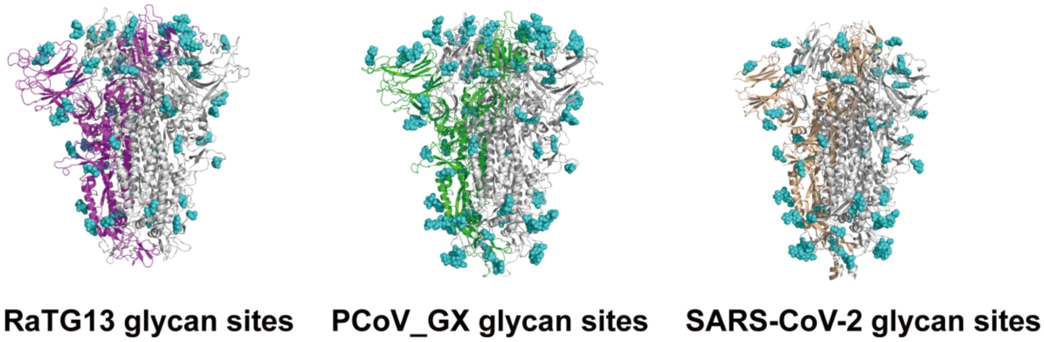


**Supplementary Fig. 4 Glycan sites of the RaTG13, PCoV_GX and SARS-CoV-2 spikes.** Glycan sites for each spike are shown, colored in cyan. A monomer for each spike trimer is colored (magenta for RaTG13 in the left panel, green for PCoV_GX in the middle panel, and wheat for SARS-CoV-2 in the right panel). The remaining monomers for each spike trimer are shown in gray.

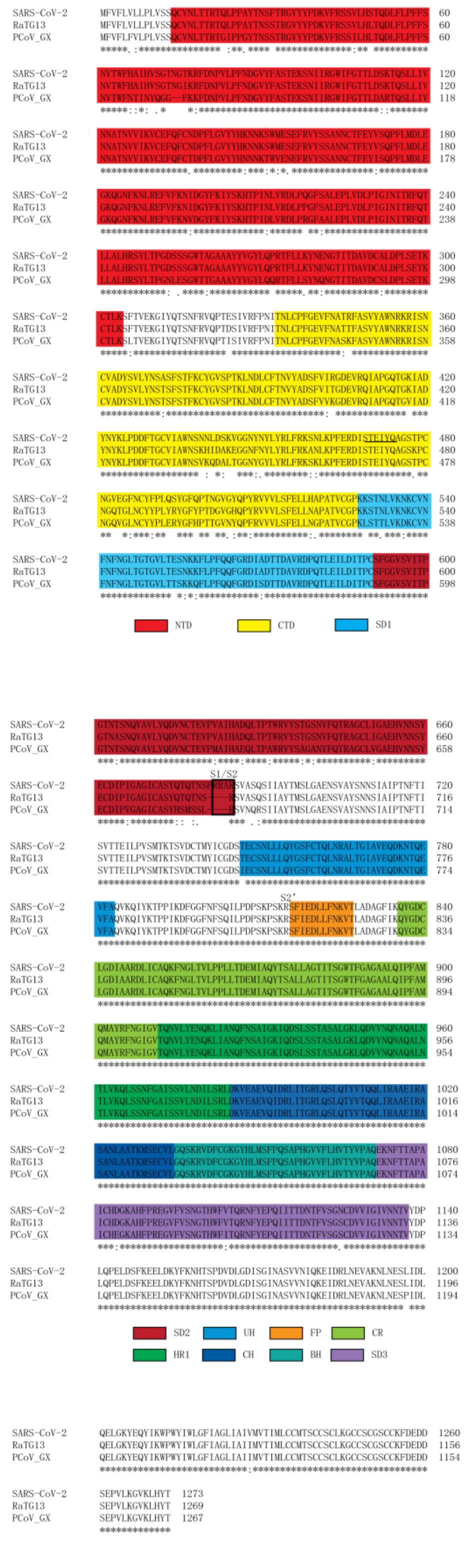


**Supplementary Fig. 5 Amino acid sequence alignment of the SARS-CoV-2, RaTG13 and PCoV_GX spikes.** Identical residues are denoted by an "*" in the bottom consensus sequence row. Structural domains are colored according to Fig. 1B.

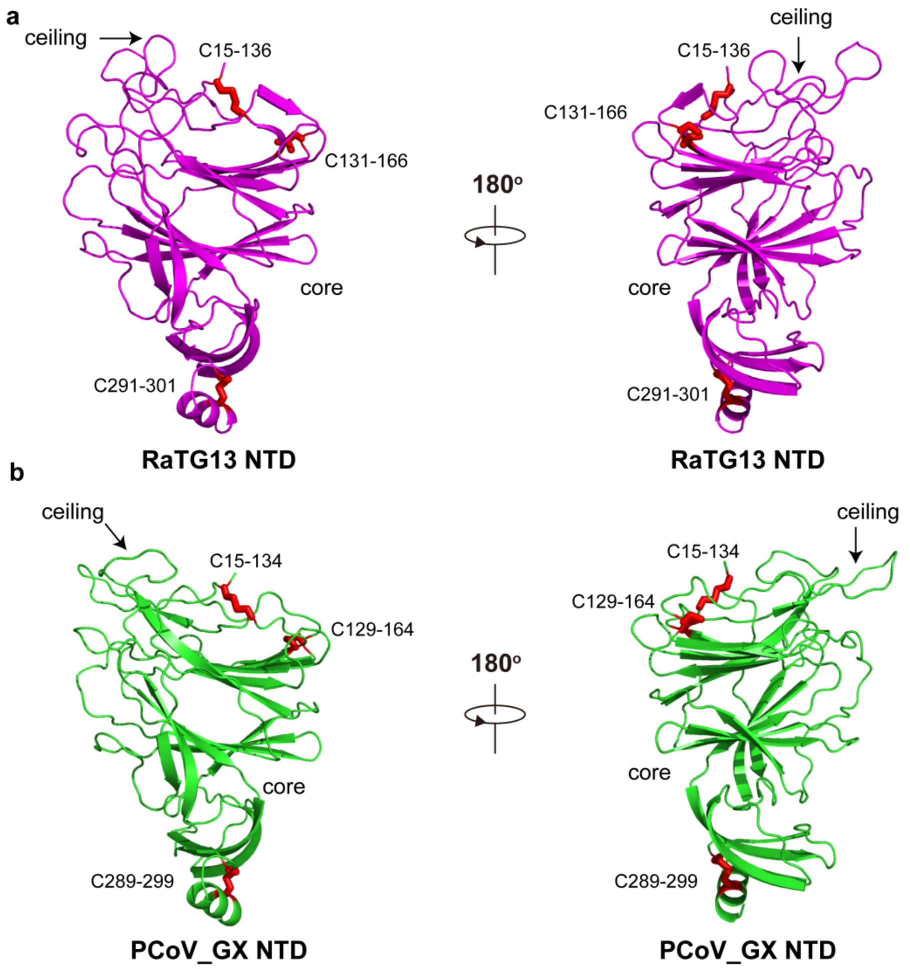


**Supplementary Fig. 6 NTD structures of the RaTG13 and PCoV_GX spikes. (a)** The RaTG13 NTD is shown in two opposite orientations and its NTD is colored magenta. **(b)** PCoV_GX NTD is shown in two orientations directions and its NTD is colored green. The ceiling region of the NTDs are indicated by black arrows, and the core regions are labeled. Disulfide bonds are shown as red sticks.

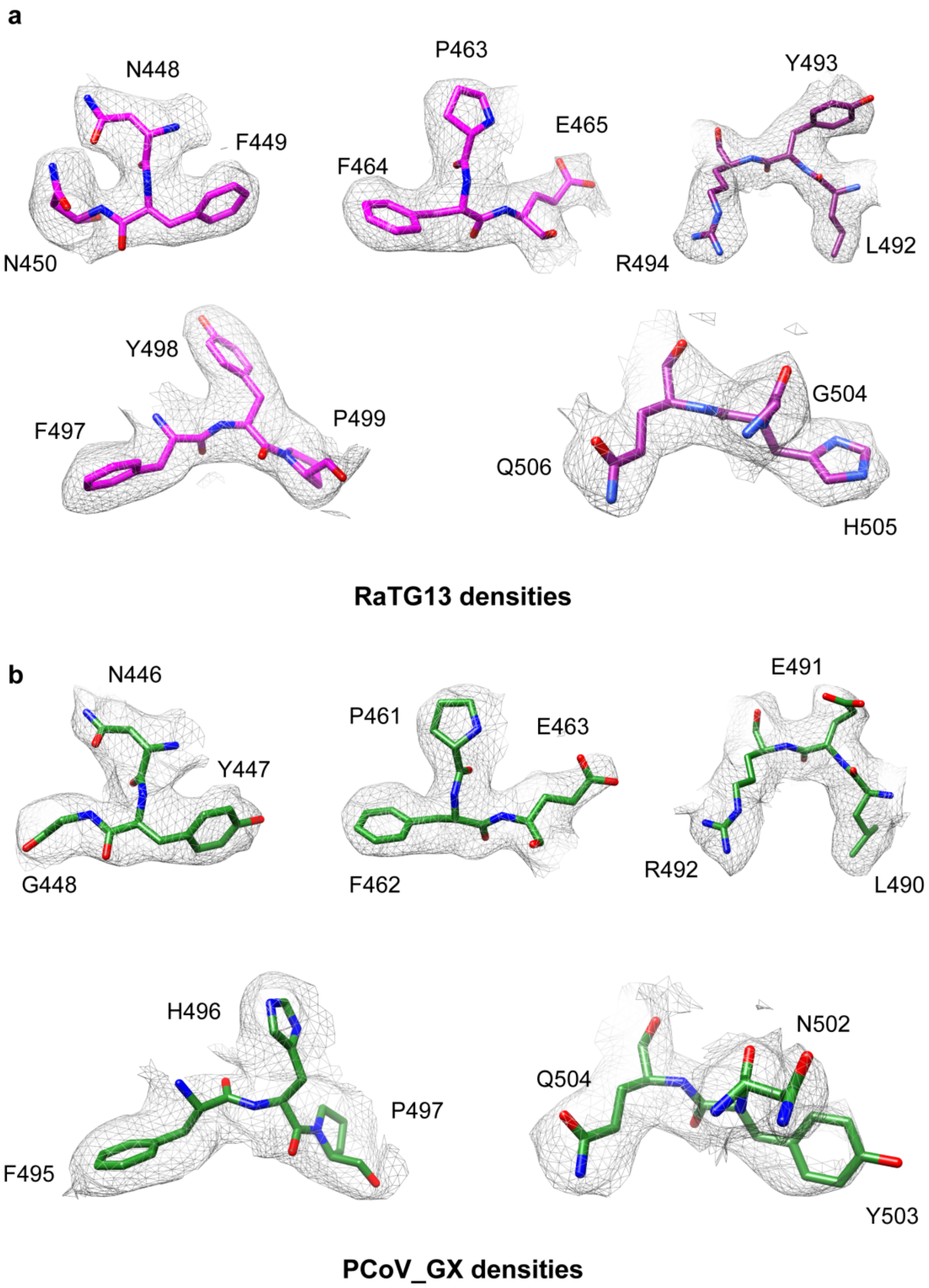


**Supplementary Figure 7. The representative density maps of the RaTG13 and PCoV_GX RBMs.** The representative density maps of the RaTG13 **(a)** and PCoV_GX **(b)** RBMs. Each map is contoured at 2.5 RMS to show the density.

**Table S1 Cryo-EM data collection, refinement, and validation statistics**

| Data collection and processing | RaTG13 (PDB-7CN4 EMDB-30416) | PCoV_GX (PDB-7CN8 EMDB-30418) |
|---|---|---|
| Magnification | ×81000 | ×81000 |
| Voltage (kV) | 300 | 300 |
| Electron exposure (e–/Å$^2$) | 50 | 50 |
| Defocus range (μm) | -1.5 to -1.8 | -1.5 to -1.8 |
| Pixel size (Å) | 1.0825 | 1.0825 |
| Symmetry imposed | C3 | C3 |
| Initial particle images (no.) | ~450,000 | ~700,000 |
| Final particle images (no.) | 99241 | 263842 |
| Map resolution (Å) | 2.93 | 2.48 |
| FSC threshold | 0.143 | 0.143 |
| Map resolution range (Å) | 2.93-6 | 2.48-5 |
| **Refinement** | | |
| Initial model used (PDB code) | 5X58 | 6VSB |
| Model resolution (Å) | 2.93 | 2.48 |
| FSC threshold | 0.143 | 0.143 |
| Model resolution range (Å) | 2.93 | 2.48 |
| Map sharpening *B* factor (Å$^2$) | -89.07 | -30 |
| Model composition | | |
| Non-hydrogen atoms | 26277 | 27477 |
| Protein residues | 3360 | 3375 |
| Ligands | 54 | 84 |
| *B* factors (Å$^2$) | | |
| Protein | 105.45 | 151.92 |
| Ligand | 99.18 | 117.83 |
| R.m.s. deviations | | |
| Bond lengths (Å) | 0.008 | 0.003 |
| Bond angles (°) | 0.776 | 0.693 |
| Validation | | |
| MolProbity score | 2.01 | 2.07 |
| Clashscore | 9.83 | 11.78 |
| Poor rotamers (%) | 0.33 | 0.1 |
| Ramachandran plot | | |
| Favored (%) | 91.68 | 92.02 |
| Allowed (%) | 8.32 | 7.9 |
| Disallowed (%) | 0.00 | 0.09 |

**Table S2 The interacting residues and glycans with one RBD of the different spikes**

| Spike | Monomer of counterclockwise | Monomer of clockwise | The number of interacting residues and glycans |
|---|---|---|---|
| **RaTG13** | 41K, 43F,113K, 115Q, 132E,167T,198D,199G, 200Y, 230P, 231I, 232G,369Y,370N, 373S, 374F, 375S, 377F, 384P, 385T, 503V,977L, 978S, 979R,980L,981D, 984E,N165-linked glycan, N234-linked glycan,N370 -linked glycan | 405D,408R,413G, 415T,416G,417K, 420D,421Y,453Y, 455L, 503V, 982P,983P, | 43 |
| **PCoV_GX** | 111R,113Q, 130E,163N, 196D,197G,198Y, 228P,229I, 230G, 232N,367Y,368N, 371S, 372F, 373S, 375F, 382P, 383T, 386N,435N,973D, 975L, 976S, 977R, 978L, 979D, N163-linked glycan,N232-linked glycan, N368-linked glycan | 401K,403D,406R, 411G,412Q,413T, 414G,418D, 451Y,453L,458K 502N,503Y,979D, 980P,981P | 46 |
| **SARS-CoV-2 (PDB :6VXX)** | 41K,200Y, 230P,369Y, 370N, 979D,981L,982S, 983R, 984L, 985D, N234 -linked glycan | 415T,416G,417K, 421Y, 987P | 17 |
| **SARS-CV-2 (PDB:6ZGE)** | 113K, 115Q, 198D, 199G,200Y,230P, 231I, 232G, 234N, 365Y, 369Y, 370N, 373S,374F, 375S, 377F, 384P, 979D, 981L, 982S, 983R, 984L, 985D, 988E,N165-linked glycan, N234-linked glycan | 403R, 405D, 408R, 413G, 415T, 416G, 417K, 420D, 421Y, 455L, 505Y, | 37 |
